# Vegetation memory effects and their association with vegetation resilience in global drylands

**DOI:** 10.1101/2021.08.22.457255

**Authors:** Erik Kusch, Richard Davy, Alistair W. R. Seddon

**Author notes:** CORRESPONDING AUTHOR: Erik Kusch;, Department of Biology, Section for Ecoinformatics & Biodiversity, Center for Biodiversity Dynamics in a Changing World (BIOCHANGE), Arhus University.

## Abstract

1. Vegetation memory describes the effect of antecedent environmental and ecological conditions on the present ecosystem state and has been proposed as an important proxy for vegetation resilience. In particular, strong vegetation-memory effects have been identified in dryland regions, but the factors underlying the spatial patterns of vegetation memory remain unknown.
2. We aim to map the components and drivers of vegetation memory in dryland regions using state-of-the-art climate reanalysis data and refined approaches to identify vegetation-memory characteristics across dryland regions worldwide.
3. Using a framework which distinguishes between intrinsic and extrinsic ecological memory, we show that: (i) intrinsic memory is a much stronger component than extrinsic memory in the majority of dryland regions; and (ii) climate reanalysis data sets change the detection of extrinsic vegetation memory effects in some global dryland regions.
4. *Synthesis*. Our study offers a global picture of the vegetation response to two climate forcing variables using satellite data, information which is potentially relevant for mapping components and properties of vegetation responses worldwide. However, the large differences in the spatial patterns in intrinsic vegetation memory in our study compared to previous analyses show the overall sensitivity of this component in particular to the initial choice of extrinsic forcing variables. As a result, we caution against using the oversimplified link between intrinsic vegetation-memory and vegetation recovery rates at large spatial scales.

## Introduction

There is a growing concern that future ecosystem disturbance dynamics will exceed ecological tipping points and cause non-linear shifts in ecosystem states. Examples of non-linear shifts in ecosystem states include regime shifts from tropical forest to savannas (Senf & Seidl, 2018), from coral-dominated to algae-dominated coral reefs (Nyström & Folke, 2001), and eutrophication and acidification processes in shallow lake ecosystems (Spears et al., 2017). Such regime shifts can have major local and global impacts with consequences for ecosystem functioning and human well-being (Côté & Darling, 2010; Genkai-Kato, 2007; Scheffer & Carpenter, 2003).

Ecological resilience, the ability of a system to withstand disturbance and maintain its general functioning, is an important property to safeguard against such regime shifts (Holling, 1973). Ecological resilience has been incorporated into the objectives of numerous agencies influential at the global policy level (e.g. Convention on Biological Diversity, 2020), and is a key focus within several of the United Nations Sustainable Development Goals (“Sustainable Development Goals,” 2020).

Although ecological resilience has been prevalent in the global change literature for at least the last three decades (Côté & Darling, 2010; Cumming et al., 2015; Foley, Coe, Scheffer, & Wang, 2003; Neubert & Caswell, 1997), universal baseline metrics on which to assess and compare resilience in different ecosystems are yet to be established (Standish et al., 2014). Consequently, it is impossible to make reliable assessments of resilience and compare ecosystem components through time and space (Pimm, Donohue, Montoya, & Loreau, 2019). Indeed, although several metrics of resilience have been proposed (i.e. resistance to perturbation, recovery rate from a disturbance, and robustness) based on ecological theory (Grafton et al., 2019; Hodgson, McDonald, & Hosken, 2015), a challenge that follows is how to operationalize these metrics with empirical data.

Remote sensing has been proposed to act as a useful tool to map components of resilience to climate variability at large spatial scales by combining data of both ecological response (e.g. vegetation indices) and forcing (e.g. climate) variables. For example, studies of vegetation responses to climate variability using remote-sensing techniques, have identified highly localised patterns of vegetation sensitivity (Seddon, Macias-Fauria, Long, Benz, & Willis, 2016), lagged-vegetation response to climatic anomalies (Liu, Zhang, Wu, Li, & Qin, 2018; Vicente-Serrano et al., 2013), vegetation resistance (an inverse proxy of how easily the system is perturbed where an easily perturbed system exhibits low resilience) and engineering resilience (a proxy for how quickly a perturbed system returns to its pre-disturbance state) (De Keersmaecker et al., 2015).

One important feature of studies which use remote sensing data to map components of ecological resilience worldwide is the quantification of vegetation-memory effects. Vegetation memory describes the effect of antecedent environmental and ecological conditions on the present ecosystem state, and can occur as a result of both ‘intrinsic’ (e.g. internal vegetation dynamics affecting the recovery rate) and ‘extrinsic’ forcing (e.g. a lagged response to climate variables) (Ogle et al., 2015). Indeed, it has been suggested that the strength of the intrinsic vegetation memory, identified using an autoregressive model in which the previous time step’s vegetation metric is used as a predictor alongside current climate variables (e.g. temperature, precipitation), can be used as a direct indicator for the recovery rate (i.e. engineering resilience). Because ecological theory indicates that increased autocorrelation occurs in models in which an ecological state variable is increasingly placed under stress and approaching a threshold (Scheffer et al., 2009), reduced recovery rate (i.e. a stronger intrinsic memory as indicated by an increased autoregressive coefficient) may be proxy of reduced ecological resilience. This approach has been used to assess changing patterns of resilience in tropical forests (Papagiannopoulou et al., 2017), Mediterranean forests (Gazol, Camarero, Sangüesa-Barreda, & Vicente-Serrano, 2018), and global drylands (De Keersmaecker et al., 2015).

However, although theoretical simulations may predict reduced recovery rates as a result of reduced ecosystem resilience, it is unclear if this simple approximation can be universally applied at a global scale. Vegetation-memory effects can occur as a result of both ‘intrinsic’ (e.g. critical slowdown resulting from reduced engineering resilience) and ‘extrinsic’ variables (e.g. a lagged response to climate variables) (Ogle et al., 2015). Whilst the first-order autoregression (AR1) coefficient may be applicable for identifying engineering resilience in some settings, this relies on the assumption that all other extrinsic variables have been accounted for. If not, then the autocorrelation coefficient used to identify intrinsic memory has the potential to mask the effect of other extrinsic forcing variables. For example, one recent study showed that some regions that exhibited high memory-effect coefficients in previous studies could be better explained by varying the length of extrinsic memory (Liu et al., 2018). As a result, the underlying applicability of this metric as proxy for ecological resilience must be considered before it can be implemented more broadly.

One region of particular relevance for such studies are global drylands. Global drylands, defined as areas in which crop production is limited by water availability (Adeel et al., 2005; Millennium Ecosystem Assessment, 2005b), are commonly observed to exhibit strong intrinsic memory coefficients, and are therefore assumed to exhibit low engineering resilience (De Keersmaecker et al., 2015). Additionally, these systems function as important carbon pools (Tian, Brandt, Liu, Rasmussen, & Fensholt, 2017) and may be prone to locally facilitated regime shifts (Xu, Van Nes, Holmgren, Kéfi, & Scheffer, 2015). Thus, bolstering the ecosystem resilience of these regions is essential in securing the reliability of agricultural landscapes for local and global communities.

Here, we investigate the factors influencing vegetation memory in global dryland regions. Specifically, we combine a state-of-the-art climate reanalysis product and satellite-based global vegetation indices to extend previous analyses of vegetation-memory effects in drylands in terms of (1) spatial and temporal coverage, (2) consistency, (3) accuracy, and (4) climatological variables at a global scale. Specifically, we investigate (i) the relative importance of extrinsic factors and intrinsic-vegetation memory for determining the vegetation response in dryland regions; (ii) the length of extrinsic memory (i.e. how long does an external climate perturbation affect the system?), and (iii) the strength of this memory (i.e. the magnitude of the vegetation memory coefficients and whether this coefficient is positive or negative). Using this information, we then consider our findings in the context of understanding patterns of ecological resilience worldwide.

## Material & Methods

Dryland regions were identified following the Millennium Ecosystem Assessment (Millennium Ecosystem Assessment, 2005b) which classifies any region as a dryland which exhibits aridity index (long-term mean ration of mean annual precipitation to mean annual evapotranspiration) values below 0.65 (Millennium Ecosystem Assessment, 2005a).

### Data Sets

We used GIMMS NDVI 3g data to identify vegetation response in global dryland regions between January 1982 and December 2015. NDVI is a compound vegetation index made up from reflectance in the red and near-infrared reflectance bands. It is highly correlated with vegetation cover (Harris, Carr, & Dash, 2014; Tian et al., 2017), productivity (Gamon et al., 1995), and photosynthetic performance (Gamon et al., 1995; Pettorelli et al., 2005). The GIMMS dataset provides bi-weekly (Stow et al., 2004) NDVI snapshots on a 0.083° × 0.083°(Liu et al., 2018) (∼ 9.27km × 9.27km) resolution at global scale. We subsequently built monthly-maximum composites to minimise the impacts of cloudy periods (Stow et al., 2004). While NDVI records tend to saturate in regions of high biomass and thus high NDVI values (Huete et al., 2002), these issues are unlikely to affect dryland regions.

For independent climate data, we used the European Centre for Medium-range Weather Forecast’s (ECMWF’s) ReAnalysis 5 (ERA5) (Balsamo et al., 2015; ECMWF, 2018a). ERA5 is created using the ECMWF’s integrated forecasting system and a large volume of satellite and ground-based data from a wide variety of data providers through the use of data assimilation (ECMWF, 2018b). ERA5 benefits from recent methodological advancements in data assimilation, incorporating a wide range and high volume of weather station observations and improved understanding of physical processes within the integrated forecasting system (ECMWF, 2018c, 2018d).

One major advantage of using climate reanalysis products to understand vegetation memory effects is that they enable the study of responses at higher temporal resolution and consistency than previous studies. For example, Liu et al., 2018, used the CRU3.0 dataset, which is a suboptimal dataset for dryland regions because it contains large areas of the globe where precipitation time series data must be replaced with mean values, thus biasing the trends (Macias-Fauria, Seddon, Benz, Long, & Willis, 2014). Reanalysis products have been widely used to shed light on physical processes (ECMWF, 2018e; Tarek, Brissette, & Arsenault, 2020) which are highly influential in determining vegetation memory properties. For our analyses, we selected two main ecosystem-drivers: soil moisture and air temperature.

**Soil Moisture** (Qsoil) was used as a proxy of local water regimes given that water-dependent vegetation memory is likely to rely on water availability to root systems (Ogle et al., 2015). Qsoil metrics were included to take into account that precipitation events may be subject to further soil processes such as pore connectivity for precipitated water to be available to plant roots (Smith et al., 2017). Therefore, we anticipated that soil moisture would serve as a more direct proxy of local water regimes compared to the drought indices used in previous studies.

ERA5 includes four distinct layers in the soil for the calculation of Qsoil indices: (1) Soil Moisture (0-7cm) (Qsoil1), (2) Soil Moisture (7-28cm) (Qsoil2), (3) Soil Moisture (28-100cm) (Qsoil3), and (4) Soil Moisture (100-255cm) (Qsoil4). Typical drought indices (e.g. SPEI) do not allow for this additional distinction. Within ERA5, unfrozen ground water 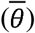 across all four soil layers (*k*) is defined as:

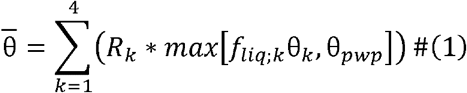

with *R*_*k*_ being the root fraction of soil layer *k* which is a fixed metric according to Land-Cover Classification Systems (LCCSs), and the statement *max*[*f*_*liq;k*_θ_*k*_, θ_*pwp*_] calculating the amount of unfrozen soil water in soil layer *k. f*_*liq;k*_ is a parametrised function of soil temperature; θ_*pwp*_ denotes the permanent wilting point according to soil texture. For a more in-depth explanation of how Qsoil is calculated within ERA5, see the IFS Documentation CY45R1 Chapter 4 Physical Processes (ECMWF, 2018e).

**Air Temperature** was used as an additional predictor in this study given its links to vegetation sensitivity (Seddon et al., 2016), tree-ring growth (Esper, Schneider, Smerdon, Schöne, & Büntgen, 2015), and global primary production (Prince & Goward, 1995), in addition to severe drought events with possible large consequences to local vegetation (Allen et al., 2010). Within this study, we used Air Temperature (at 2m above ground) (Tair) as contained within the ERA5 data set, due to the demonstrated impact of Tair on different aspects of plant physiology and plant morphology which may manifest in vegetation memory effects.

ERA5 data is available for hourly intervals (which we averaged to monthly-means) from 1950 to present day at a 30km × 30km spatial resolution of global coverage making the resolution of ERA5 and AVHRR-based GIMMS NDVI 3g incompatible. We resolved this issue by statistically downscaling ERA5 data using kriging (see Supplementary Information: “Statistical Downscaling”).

### Statistical Approach

To assess the relative importance of intrinsic memory and extrinsic climate forcing we used a linear modelling approach (e.g. De Keersmaecker et al., 2015). Our vegetation memory models are based on the following specification:

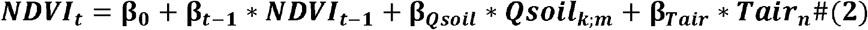

with *NDVI*_*t*_ and the Autoregressive NDVI Coefficient (*NDV I*_*t*−*1*_) being standardised NDVI anomalies at time step *t* and *t* − 1, respectively; *Qsoil*_*k;m*_ denoting Qsoil data at depth level *k* (translating to Qsoil1-Qsoil4) and cumulative lag of standardised anomalies of lag *m*, and *Tair*_*n*_ denoting Tair data as the cumulative lag of standardised anomalies of lag *n*. We extracted data for each variable (NDVI, Tair, Qsoil) and detrended it using linear detrending to avoid effects of changing abiotic conditions over long time-series (De Keersmaecker et al., 2015). Subsequently, we standardised the detrended data to Z-Scores to obtain deviations of monthly means/monthly anomalies for each variable (De Keersmaecker et al., 2015):

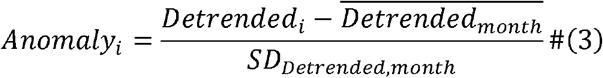

with *i* indexing individual, detrended data records. This resulted in a final set of monthly time-series of vegetation and environmental dynamics expressed as divergences from expected seasonal trends across the study regions. Lagged effects were calculated as (1) *NDVI*_*t*−*1*_ (to identify intrinsic vegetation memory) using Z-Score NDVI data, and (2) cumulative lags of detrended *Qsoil/Tair* data (to identify extrinsic Qsoil/Tair-driven memory) with lags ranging from 0 (instantaneous effects) to annual effects (aggregated over twelve months of detrended Qsoil/Tair records) in steps of one month at a time. These lagged effects are subsequently standardised to Z-scores.

We used Principal Component Analysis (PCA) regression to limit the effects of collinearity in our forcing variables. We did so for each of the cumulative *Qsoil* lags across all four Qsoil layers. Firstly, z-score data for *NDVI*_*t*−*1*_, *Tair*, and *Qsoil* are analysed using a PCA. Secondly, linear regression was performed as follows:

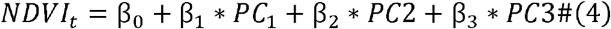

with *NDVI*_*t*_ representing NDVI anomalies; these were set to NA (skipped in models) in months for which *Thresholds*_*i*_ < 0.1 with 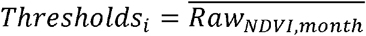 to circumvent unreliability of NDVI data at low scores (Liu et al., 2018). PC1 through PC3 and β_l_ through β_3_ indicate principal components 1 through 3 and coefficients of their effects in the model respectively. Thirdly, we performed model selection to identify the cumulative soil moisture and air temperature lags which present the most explanatory power through comparison of the model Akaike Information Criterion (AIC) values (lowest AIC indicates best model performance). This results in a proxy of Qsoil- and Tair-memory length of local vegetation. Finally, the regression coefficients *β*_l_ through *β*_3_ can be back-transformed to represent PCA input variable effects (see formula 1) using PCA loadings and PCA model coefficients. See the Supplementary Information: “Model Workflow and Interpretation” for a visual representation of the workflow and an overview of model coefficient interpretation.

#### Model Comparisons

In order to determine which vegetation memory component (i.e. model variable) has the greatest influence on vegetation anomalies, we first compared the absolute values of variable coefficients across all pixels within each model individually, separately for a set of five pre-selected study regions (i.e. South-Eastern Europe, the Caatinga (N-E Brazil), Australia, the contiguous US, and the Sahel region) which exhibited marked patterns of strong vegetation memory in contemporary studies (De Keersmaecker et al., 2017, 2015; Liu et al., 2018; Seddon et al., 2016; Vicente-Serrano et al., 2013). This was done to reduce computational requirements in the downscaling process (i.e. downscaling all four Qsoil levels for our pre-selected study regions and only the most biologically important Qsoil layer globally). Additionally, to identify which Qsoil layer is the most biologically influential, we compared absolute values of *Qsoil* coefficients for all pixels per model between all four models for each of these five pre-selected study regions using Mann-Whitney U-Tests. Across all pre-selected study regions (South-West Europe, Australia, Caatinga, the contiguous US, and the Sahel region), *Qsoil1* (soil moisture in a band of 0-7cm depth) was identified as the most meaningful out of all the soil moisture variables. Therefore, we carried out the global analysis presented going forward using only *Qsoil1* input out of the four available soil moisture parameters within ERA5.

#### Variance Partitioning

To assess the relative importance of intrinsic and extrinsic vegetation memory components in driving the system’s response to disturbances, we used variance partitioning on each pixel in our study. In this procedure, we determined the relative importance contained in the vegetation memory model predictors *NDVI*_*t*−*1*_ (intrinsic memory), *Qsoil*_*k;m*_ (extrinsic soil moisture memory of layer *k* and cumulative lag *m*), and *Tair*_*n*_ (extrinsic air temperature memory at cumulative lag *n*) using a partial regression approach (see Supplementary Information: “Variance Partitioning”). This allowed us to map the relative importance of components of vegetation memory, and spatial inspection of which variables best explain NDVI anomalies.

## Results

### Vegetation Memory Properties

According to AIC scores, our vegetation memory models performed best across parts of Saudi-Arabia, the Mongolian steppe, and parts of Australia (Figure 1). Global dryland regions are characterised by strong, positive intrinsic vegetation-memory effects, with the strongest values across Australia, South Africa, Texas, and Algeria/Morocco and weakest across the Sahara Desert, and the Russian steppe.

**FIGURE 1.**
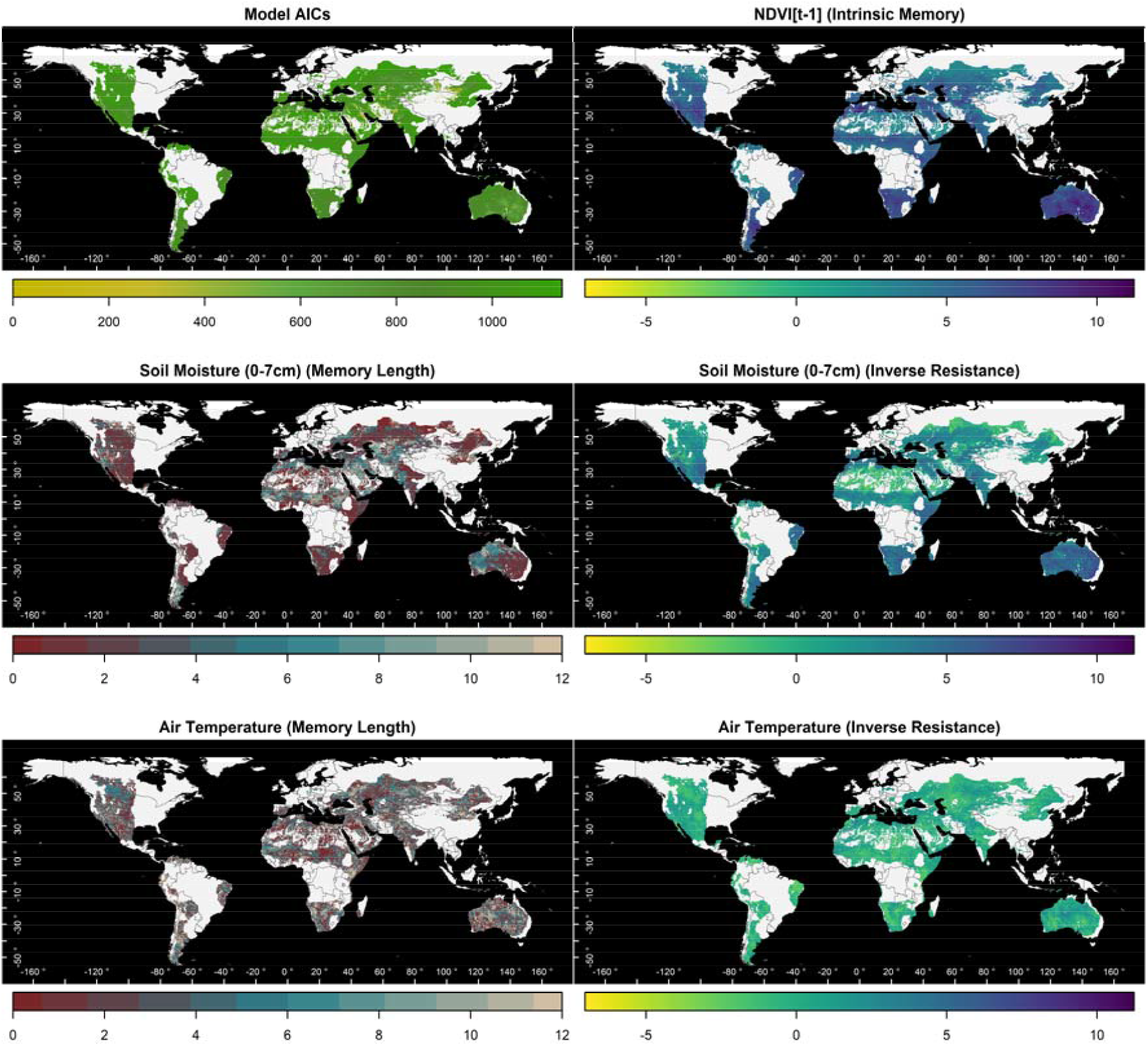
VEGETATION MEMORY PROPERTIES. – Global drylands show highly localised patterns of vegetation-memory model components. Regions of low AICs (i.e. good model fit) coincide with areas of high intrinsic memory. Patterns of intrinsic vegetation memory are mirrored by patterns of soil-moisture memory effects. Negative effects of soil-moisture memory are linked to instantaneous soil-moisture memory responses. Air temperature memory effects are negative across almost all of the global dryland regions. Note that colour palette between all outputs pertaining to the same vegetation memory characteristic (i.e. strength or length of memory) have been synchronized to make the outputs of regions comparable. For an interactive visualisation of these results, please use: https://erikkusch.com/wp-content/uploads/2020/09/GlobalDrylands_Effects.html.

The patterns of intrinsic vegetation memory are mostly mirrored by soil-moisture driven vegetation-memory effects, which exhibit strong, positive values across the dryland areas (i.e. dryland-vegetation performance is enhanced by surplus soil moisture). The positive effects of soil-moisture memory are most prevalent across Australia, South Africa, west of the Gulf of Mexico, the Horn of Africa, and the Caatinga. However, some negative soil-moisture memory (i.e. adverse effects of surplus water anomalies on local NDVI values) were also identified, in areas of low intrinsic-vegetation memory such as the northern regions of drylands in the Russian steppe, the Sahara desert, north-eastern China, south Saudi-Arabia, and parts of the south-western contiguous United States.

Extrinsic soil-moisture memory lags range from fast-responding vegetation communities (i.e. soil-moisture memory length close to 0 months) to regions of slow-response (i.e. soil-moisture memory length close to 12 months). While clear global patterns of soil-moisture memory lags are difficult to establish in general, we find a clear west-east gradient of soil-moisture memory length across both Australia and North America. Additionally, soil-moisture memory length in Eurasia and Africa increases from north to south. Finally, areas of negative soil-moisture memory effects are almost exclusively characterised by soil-moisture memory lags of zero months (i.e. instantaneous responses).

Air-temperature-driven memory characteristics show strong, negative effects across many dryland regions (i.e. dryland-vegetation performance is diminished by warmer temperatures), but are not limited to negative responses globally. These effects are strongest across South Africa, the Caatinga, parts of Australia, and Kazakhstan. However, there are areas where positive air-temperature memory effects can be observed. These indicate an increase in local vegetation productivity following/coinciding with positive air-temperature anomalies and can be observed across Algeria, parts of the Mongolian steppe, parts of Australia as well as Polish and Turkish drylands. Air-temperature memory lags manifest as highly localised patterns of vegetation response time to air temperature anomalies.

### Relative importance of vegetation-memory components

Although there are clear spatial patterns in the different components of vegetation memory, in general, the relative strength of the intrinsic (i.e. AR1) variable is much greater than the extrinsic variables. Overall, vegetation-memory effects are strongest for intrinsic vegetation memory, followed by soil-moisture memory, and air-temperature memory thus identifying a hierarchy of importance of these parameters in shaping the vegetation performance/NDVI values of local dryland vegetation through time and space (Figure 2). Here, intrinsic vegetation memory (*NDVI [t-1]*) and soil moisture effects control most of the global dryland vegetation performance in our vegetation memory modelling approach.

**FIGURE 2.**
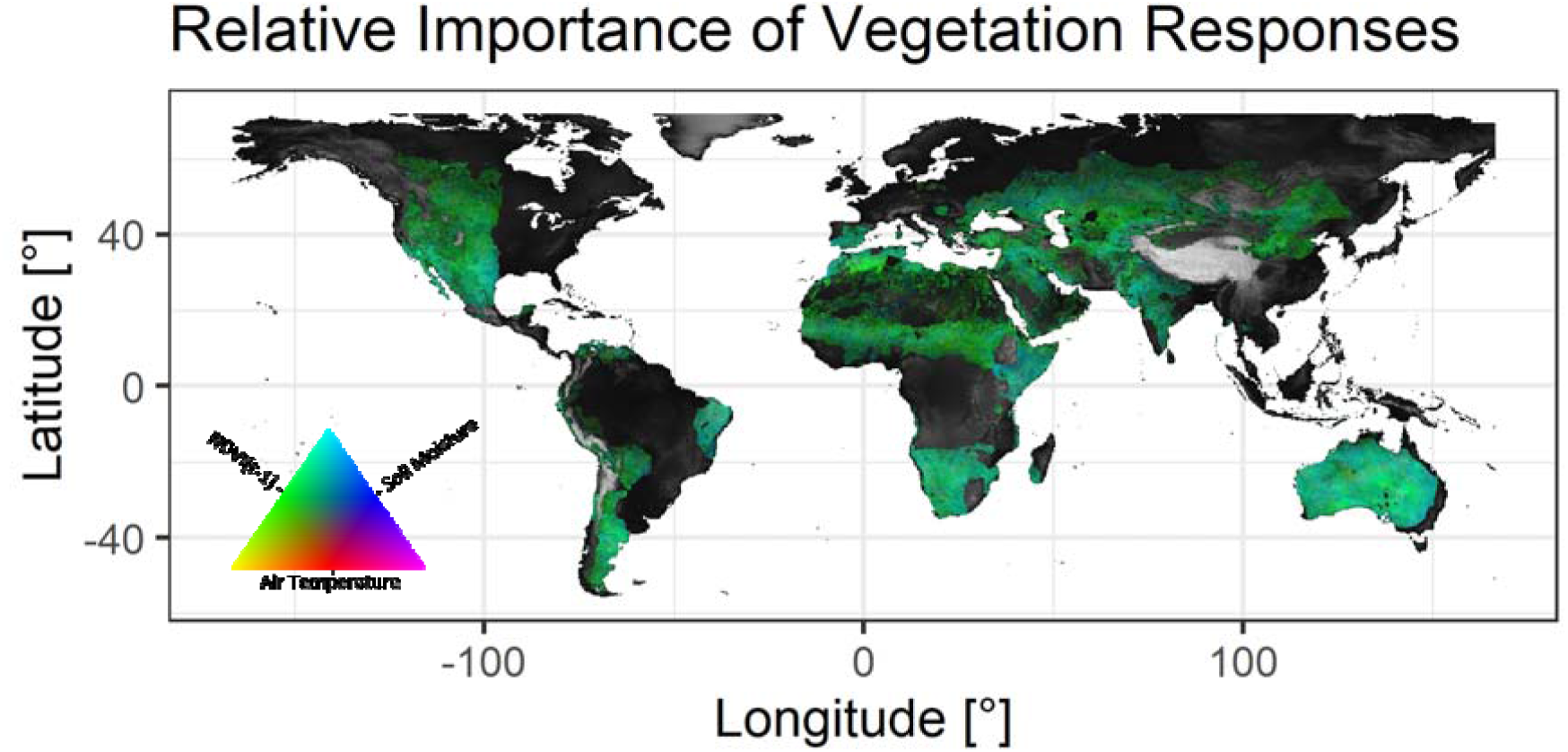
VEGETATION MEMORY EFFECT MAGNITUDE. – Average contributions of vegetation-memory effects to the local vegetation performance. To highlight only the strength of effects, model coefficients have been converted into absolute values.

Similarly, although patterns of intrinsic and extrinsic vegetation memory components are strikingly similar in terms of their spatial patterns (see Figure 1), variance partitioning indicates the majority of information is contained within intrinsic vegetation memory effects (*NDVI*_*t-1*_) with *Qsoil1* memory and shared information between the two taking up almost all of the remaining explained variance (Figure 3).

**FIGURE 3.**
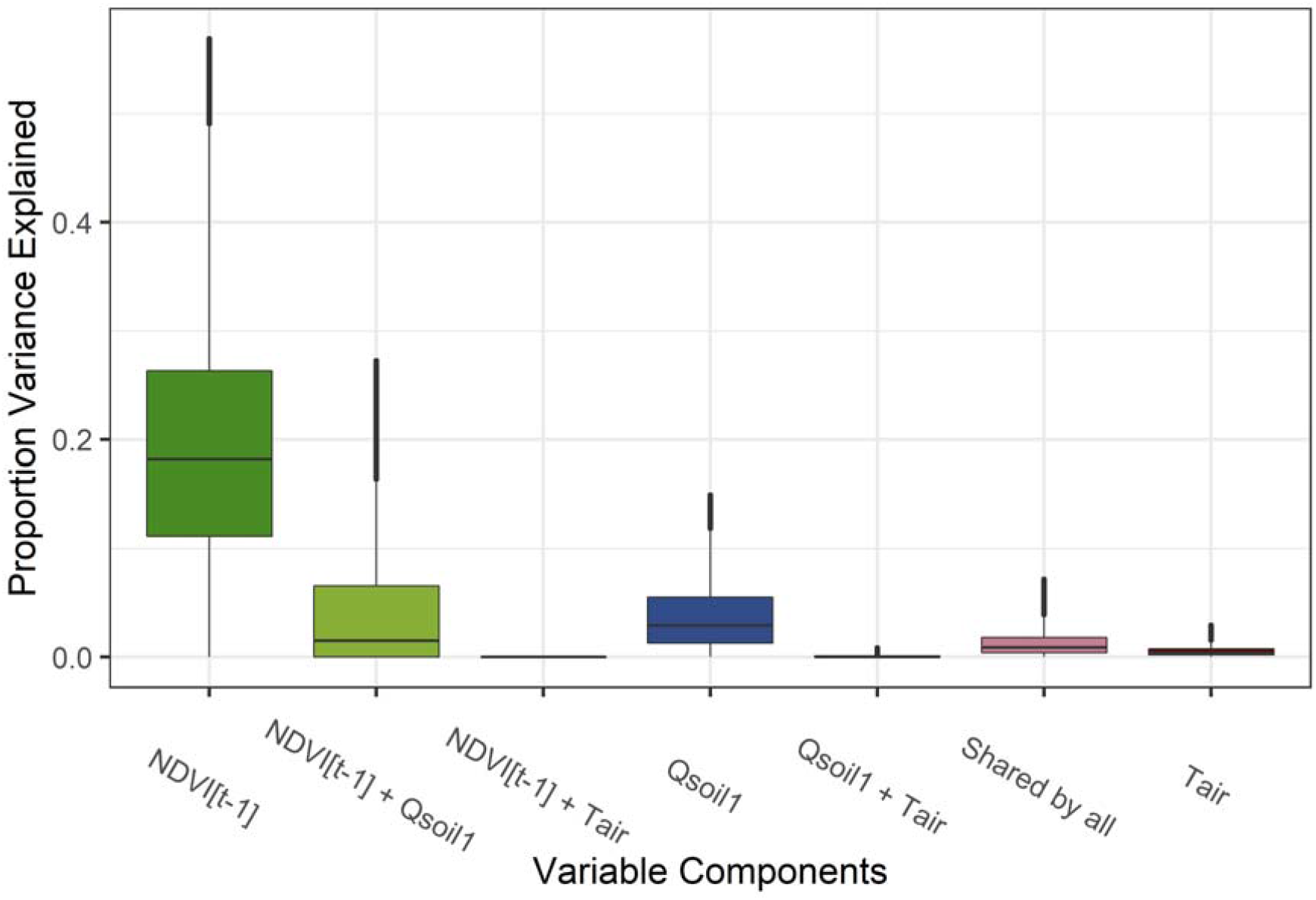
VEGETATION MEMORY VARIANCE PARTITIONING. – Variance in anomalies of NDVI explained by model parameter across global dryland raster cells. Variances portrayed here have been limited to 95% quantiles of their respective ranges.

As such, almost all explanatory power of our vegetation memory models is contained within NDVI_*t-1*_ and Qsoil1 data making these the two prominent factors driving vegetation memory in global drylands. However, the comparatively high values of variance shared between these two variables shows that our models can find it difficult to distinguish intrinsic from extrinsic vegetation memory (Supplementary Information: Figure 8 and Figure 9). This is also confirmed by the clear overlap of spatial patterns (Figure 1).

## Discussion

Our vegetation-memory approach builds on previous studies which have identified components of vegetation memory using satellite data (e.g. De Keersmaecker et al., 2015; Liu et al., 2018; Seddon et al., 2016; Vicente-Serrano et al., 2013), by using state-of-the-art climate reanalysis data (ERA5) combined with statistical downscaling. In addition, we attempted to isolate the effects of intrinsic and extrinsic memory processes for global drylands worldwide.

### Patterns of vegetation memory in global drylands

One key observation from this study is that the *NDVI*_*t-1*_ coefficient dominates in terms of the variance explained in our models across large parts of the Earth’s surface (Supplementary Information: Figure 8 and Figure 9). This result occurred in spite of the fact that our models incorporated the use of climate reanalysis data. One interpretation from these results is that, overall, global-dryland regions are characterized by strong intrinsic vegetation memory, and that this effect is – in general - much stronger than any extrinsic-memory effect identified as a result of ERA5 soil moisture or temperature variability (Figure 2 and Figure 3).

A second key finding of our study is represented by the strength of extrinsic vegetation-memory effects which vary considerably across regions, although never to the same magnitude as the intrinsic *NDVI*_*t-1*_ coefficient. We found that soil-moisture memory exerts a much greater effect on dryland vegetation than air temperature, with global-dryland vegetation to responding positively to wetter conditions (i.e. positive *Qsoil1*) and negatively to neutral to elevated air temperatures (i.e. *Tair*), suggesting that plants in these regions operate at or above their temperature optimum and below or at their soil moisture optimum.

Counterintuitively, our study identified several areas in which positive soil-moisture anomalies lead to a decrease in vegetation productivity (negative NDVI anomalies). These effects coincide with response times of one month and faster. We interpret this to be a sign of positive soil-moisture anomalies in these regions to be coupled with severe downpour and flooding events which can lead to an initial decrease in vegetation performance in certain vegetation communities (Broich, Tulbure, Verbesselt, Xin, & Wearne, 2018; Thapa, Thoms, & Parsons, 2016). Note, however, that the negative coefficients of soil-moisture memory are far smaller in extent and magnitude than their positive counterparts. Additionally, our analysis revealed that global dryland vegetation performance can react to positive air temperature anomalies in one of two ways: (1) negatively at different time lags, or (2) positively at long time lags. This may be indicative of different physiological strategies and capabilities of dealing with increased heat-stress (Wang, Heckathorn, Mainali, & Tripathee, 2016).

Patterns of extrinsic-memory length also vary considerably within and between regions. One example of this is the east-west gradient in soil-moisture memory length across Australia, with shorter responses in the east and longer responses in the west. Other regions exerting short vegetation memory to soil-moisture anomalies are located in south and east Africa as well as the Caatinga and parts of Argentina while longer soil-moisture memory is especially present throughout the Sahel region. Much like with extrinsic-memory strength, these patterns are likely a consequence factors including (but not limited to) (1) local antecedent disturbance regimes (Yin, Li, Huang, Si, & Bai, 2015), (2) plant function (Nielsen, James, & Drenovsky, 2019), (3) life history strategies (Adier et al., 2014; Archibald, Hempson, & Lehmann, 2019), and (4) soil properties (Smith et al., 2017; Wan et al., 2016). Further studies of these regions at higher spatial resolutions which meaningfully link vegetation plot data to remote sensed data are needed to uncover the underlying processes that enable this vegetation memory behaviour.

### The use of ERA-5 data in vegetation response studies

One major advantage of using climate-reanalysis products to understand vegetation memory effects is that they enable the study of responses across temporal periods of finer resolution (which our study could not make use of due to the NDVI data being used at monthly intervals) and with a greater spatial coverage compared to previous studies. For example, Liu et al., 2018, used the CRU3.0 dataset, which provides a gridded dataset based on global weather station data for air temperature, precipitation and solar radiation. However, because large areas of the Earth’s surface do not contain weather stations providing consistent time-series, many areas (particularly in tropical regions), in many cases precipitation values are replaced with climatological mean values (Macias-Fauria et al., 2014). This is a problem for analyses based on understanding patterns ecological resilience in dryland regions, since the vegetation response to climate variability is likely to be an important component, and a consequence of this is that large areas of the Earth’s surface are required to be masked from any analysis. The ERA5 data allowed to us to provide an assessment of three vegetation-memory components for dryland ecosystems worldwide without encountering these issues of temporal gaps in the data (ECMWF, 2018e).

A second advantage is that the ERA5 data enabled us to incorporate the use of soil-moisture as a bioclimatic variable to reflect water availability in dryland regions. Our expectation was that soil moisture would be a better predictor than, for example, the weather station inferred SPEI drought index because 1-3 month lagged precipitation indices have been shown to be a predictor of vegetation productivity in dryland regions as a result of soil infiltration processes. Since soil moisture availability would represent a more direct indicator of the water available to the plant, we expected this to be a strong predictor in dryland regions.

However, despite the incorporation of soil moisture data in our analysis, the autoregressive parameter (i.e., the indicator of intrinsic memory) remained the strongest predictor across large areas of the Earth’s surface, and in some cases the relative strength of the coefficient for soil moisture was actually less than that when drought indices have been used (e.g. de Keersmaecker et al. 2015). Furthermore, the patterns of soil-moisture memory lags we identified do not coincide with the precipitation-driven memory lag identified by Liu et al., (2018) or the SPEI-informed memory lag by Vicente-Serrano et al., (2013). Such variations in the results between studies suggest that ground-based validation is required to understand better the differences in these approaches of the different datasets, and to determine whether ERA5 data provide a useful tool for future research for assessing global patterns in vegetation dynamics in response to climate variability.

### The use of Intrinsic Memory as an indicator of engineering resilience

Intrinsic vegetation memory, usually identified as autoregressive coefficients of vegetation indices (e.g. Seddon et al., 2016), is increasingly being used as an indicator of vegetation engineering resilience (i.e. recovery speed) (De Keersmaecker et al., 2015; Liu et al., 2018). Our analysis, which involved using ERA-5 data to assess the factors influencing vegetation memory in global drylands reveals some important caveats to this assumption for two main reasons.

Firstly, the spatial patterns demonstrating the relative strength of the autoregressive parameter (i.e. intrinsic vegetation memory e.g. *NDVI*_*t-1*_) vary considerably across the various studies which have quantified the value of intrinsic vegetation memory relative to different extrinsic vegetation-memory variables. For example, our results revealed relatively high levels of intrinsic vegetation memory across Australia, a weak west-to-east gradient of increasing correlation between NDVI anomalies and soil moisture anomalies, and a strong west-to-east gradient of response time to soil-moisture anomalies (see Figure 4). These results are similar to Liu et al., 2018, but are in contrast to those by Seddon et al., 2016 and De Keersmaecker et al., 2015, who identified a clear west-to-east gradient of intrinsic vegetation memory (AR1 coefficient) and extrinsic drought-driven memory, respectively. Whilst the model specifications are similar in these studies, the main differences are in the choice of datasets used to represent extrinsic forcing. For example, the model in Liu et al., 2018 did not incorporate any extrinsic forcing variable, whilst Seddon et al., 2016 and De Keersmaecker et al., 2015 used a combination of satellite derived vegetation and water availability/drought indices (Figure 4b,c).

**FIGURE 4.**
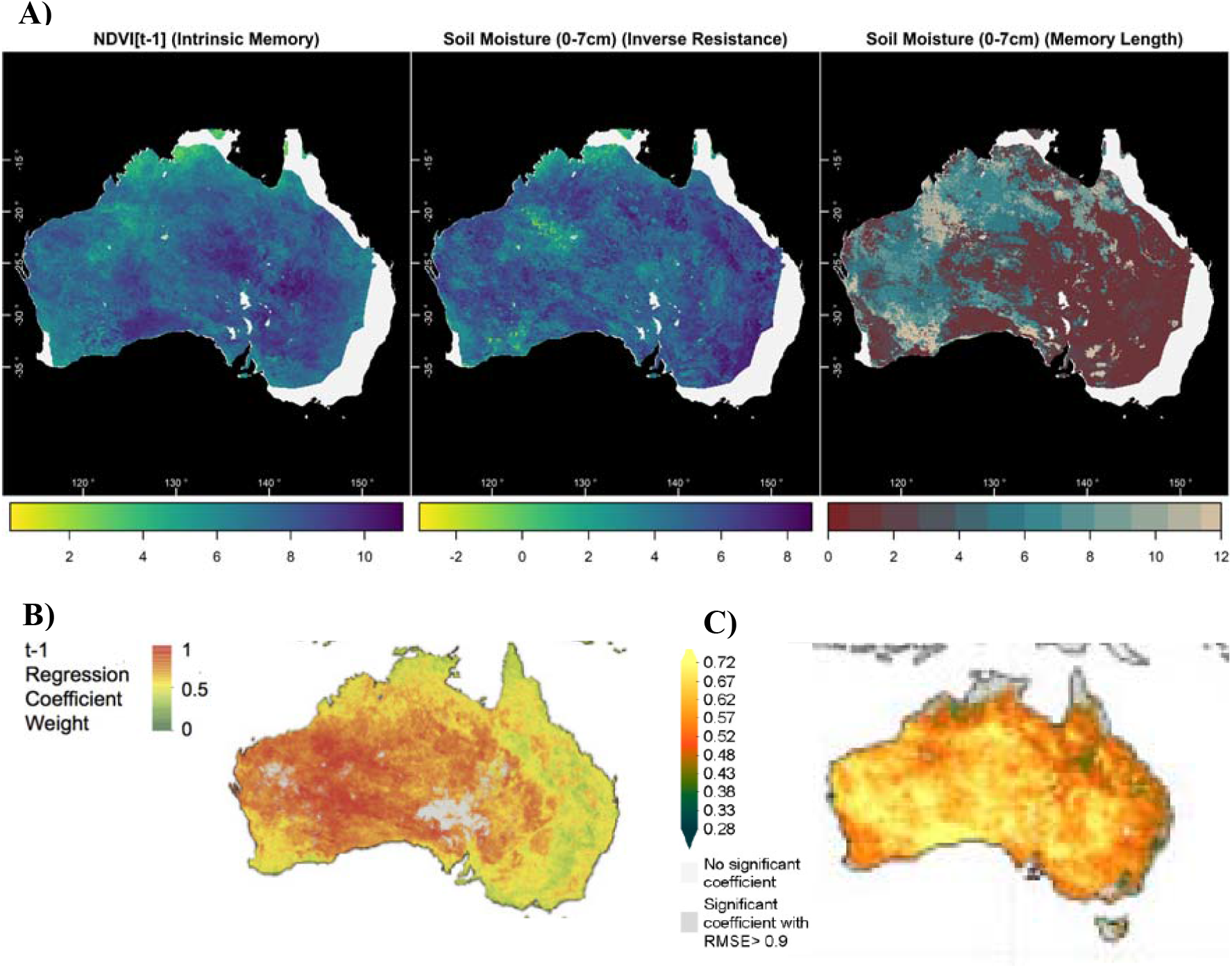
PATTERNS OF VEGETATION MEMORY COMPONENTS ACROSS AUSTRALIA -. Comparison of vegetation memory model components sourced from different contemporary approaches across Australia. (A) This study shows homogenous *NDVI*_*t-1*_ and diffuse extrinsic soil-moisture vegetation-memory patterns as well as a clear west-to-east gradient in soil-moisture-memory length which matches the west-to-east gradients of intrinsic vegetation memory by (B) Seddon et al., 2016, and (C) De Keersmaecker et al., 2015.

Similar findings are observed across the Caatinga region in NE Brazil. Here, Seddon et al., 2016 found relatively low intrinsic memory compared to the response in water availability, whilst our study and De Keersmaecker et al., 2015 found intrinsic memory to have a higher overall importance than soil moisture in this region.

These examples indicate an important caveat with regards to interpreting spatial patterns in the NDVI_*t-1*_ coefficient across different analyses, namely, that the relative importance of this variable in any model will vary strongly depending on the input variables used. As a result, interpretations of components of vegetation resistance and recovery at large spatial scales may be highly sensitive to the original forcing variables used. For example, in previous studies, eastern Australia and the Caatinga have been interpreted as being resistant to precipitation anomalies, whilst the high autocorrelation coefficient in western Australia may be interpreted as having an overall lower vegetation resilience (e.g. De Keersmaecker et al., 2015). In our study the west-east gradient in this autoregressive parameter does not exist to the same extent, and different interpretations arise from the inclusion of slightly different variables.

This issue is further compounded by temporal aggregations of input variables. By enabling the selection of the optimum soil-moisture memory-length (i.e. between 0-12 months), our model had a more flexible approach to the identification of the strength of an extrinsic forcing variable compared to those using only a fixed time window. This increase in model flexibility may also have resulted in different patterns of intrinsic memory being observed. In the Australia example, a west-east gradient is observed in extrinsic memory length to soil moisture anomalies in our study (Figure 4a, right panel), a pattern which resembles the west-east gradient in *intrinsic* memory identified Seddon et al., 2016 and De Keersmaecker et al., 2015 (Figure 4b,c). Thus, again, these results suggest a compensation effect of the AR1 coefficient in model when a fixed (e.g. 1- or 3-month lag) vs. flexible (i.e. 0-12-month lag) extrinsic variable is used.

## Conclusion

Our vegetation memory analysis using state-of-the-art climate reanalysis data, novel climatic variables and modelling methodology (in the field of vegetation memory) revealed patterns of intrinsic and extrinsic vegetation memory across global dryland regions. However, in spite of incorporation of ERA5 data to reveal patterns of vegetation memory at a global scale, we found major differences in the relative strength of intrinsic vegetation memory compared to previous studies. As a result, our study reveals the sensitivity of this variable to the incorporation of other extrinsic forcing variables, and the current understanding of intrinsic vegetation memory as a proxy of engineering resilience/recovery speed may be an oversimplification. A critical next step will be to develop a more refined selection of variables and localised identification of vegetation memory processes.

## Authors’ contributions

E.K and A.W.R.S. developed the framework used in this study. E.K. created all R scripts necessary for analyses with input by AW.R.S. R.D. created scripts for downloading ERA5 data and introduced statistical downscaling to the methodology. E.K. led analysis with contributions from AW.R.S. and R.D. All authors contributed critically to the drafts and gave final approval for publication.

## Data availability

All data sources are open access and the analysis is fully reproducible using the scripts provided at https://github.com/ErikKusch/Vegetation-Memory.

