## Supplement for "Vegetation memory effects and their association with vegetation resilience in global drylands"

### Supplementary Information

Statistical Downscaling

In order to calculate vegetation memory effects, vegetation data and abiotic data need to be represented at the same spatial and temporal resolution. Whilst both NDVI and ERA5 data are available at monthly intervals (after pre-processing as described), their spatial resolutions differ (e.g. 9.27km× 9.27km, vs. 30km × 30km, respectively). There are two ways of remedying this spatial mismatch: (1) aggregating GIMMS data to the coarser resolution of ERA5 data, or (2) downscaling of ERA5 data to GIMMS resolution. Whilst aggregating data to coarser resolutions is much easier and less computationally expensive, we opted to apply downscaling to the ERA5 data to retain valuable information within the GIMMS NDVI 3g data set. This allowed for more precise vegetation memory identification.

We used Kriging – a statistical downscaling method that is well-understood and has long been used in non-biological sciences for geostatistical interpolation purposes (Hengl, 2011). Kriging is a two-step process. First, one establishes statistical relationships between data which is to be kriged at its native resolution with covariate data at the same resolution. The second step sees the extrapolation of these relationships using covariate data at the target resolution. The way in which Kriging marks an improvement over other statistical downscaling methods lies in the fact that the Kriging methodology not only extrapolates relationships but residuals as well (see Figure 5 for a visual representation).

Therefore, Kriging within our analyses requires three inputs: (1) ERA5 data at native resolution, (2) Harmonized World Soil Database (HWSD) Digital Elevation Model (DEM) covariates at ERA5 resolution, and (3) HWSD DEM covariates at GIMMS resolution to produce sets of ERA5 data at GIMMS resolution. Kriging operations are built around formulae which establish response and predictor relationships. Our kriging approaches are specified as follows:

$$\begin{aligned} Var_{ERA5}=\alpha+\sum_{i=1}^{14} \left( \beta_{i}*Cov_{HWSD;i} \right)\#\left( 5 \right) \end{aligned}$$

with $Var_{ERA5}$ identifying any of the five ERA5 variables Tair, Qsoil1, Qsoil2, Qsoil3, or Qsoil4 at any given monthly interval between January 1981 and December 2015. $Cov_{HWSD;i}$ indexes the $i$’th HWSD covariate ranging from elevation and slope aspects to slope incline levels. Due to computational expense, the kriging procedure only considers interaction effects between slope incline levels and slope aspects (not shown in formula 5).

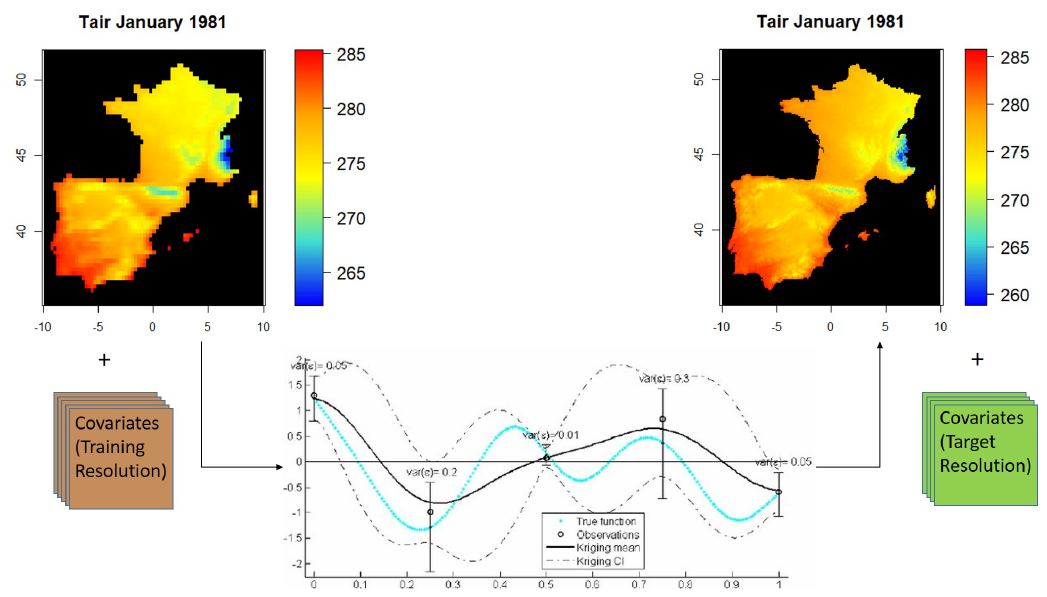

Figure 5 - Kriging Concept - Statistical downscaling effects of Tair for the time step of January 1981 using HWSD covariate data across the Iberian Region. Diagram source: Le Riche et al., 2012

For an in-depth mathematical explanation of the Kriging methodology, see Hengl, 2011. A practical example of its use can be retrieved in Lichtenstern, 2013.

Model Workflow and Interpretation

Workflow

Figure 6 presents a visual representation of the vegetation memory modelling procedure described in the publication whilst Figure 7 provides a visualisation of transformation of PCA regression model coefficients to vegetation memory model coefficients following the methodology presented by Zuur et al., 2007.

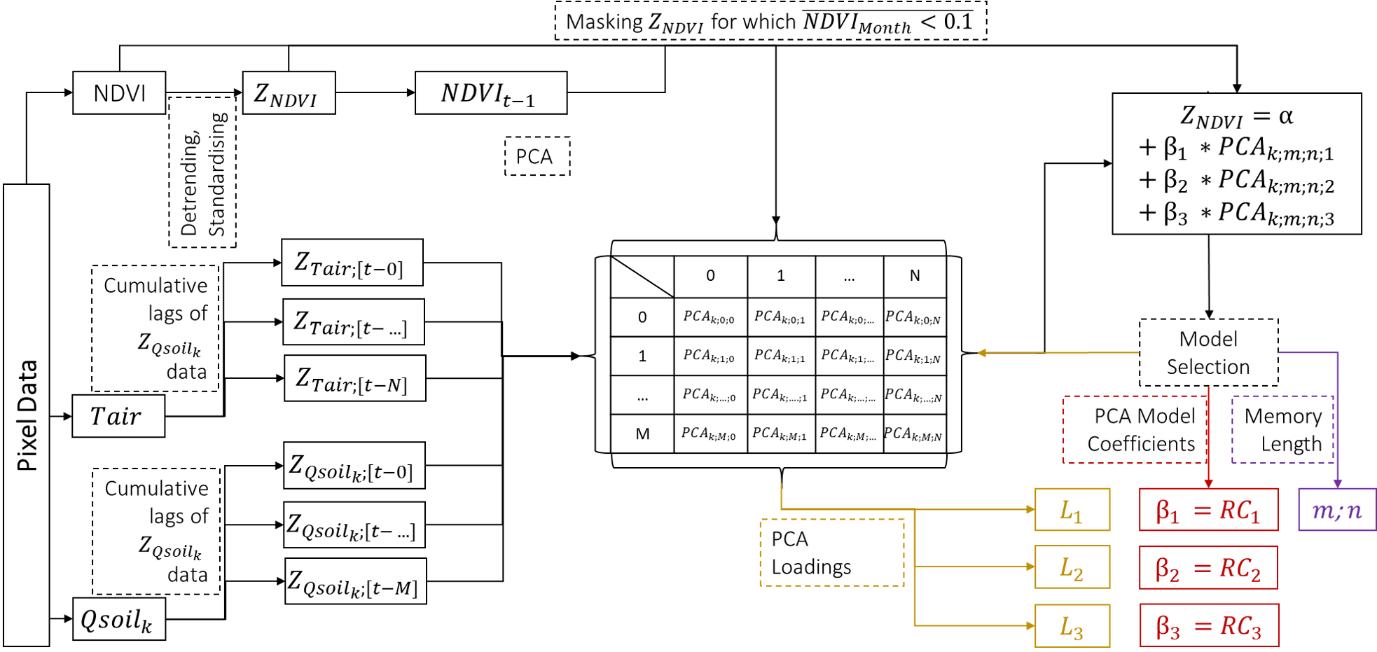

Figure 6 - Vegetation Memory Model Flowchart - Visual workflow of pixel-wise iterated vegetation memory model. $\boldsymbol{m}$ and $\boldsymbol{n}$ denote the currently considered cumulative lags of Qsoil and Tair data with $\boldsymbol{M}$and $\boldsymbol{N}$ being the maximum cumulative lag. Qsoil layers Qsoil1- Qsoil4 are identified via $\boldsymbol{k}$. $\boldsymbol{L}_{\boldsymbol{1}}$ through $\boldsymbol{L}_{\boldsymbol{3}}$ are the loadings of each detrended and standardised variable (NDVI, Tair, and Qsoil) onto the principal components 1 through 3, respectively. PCA model coefficients are identified as $\boldsymbol{\beta}_{\boldsymbol{1}}$ through $\boldsymbol{\beta}_{\boldsymbol{3}}$. Finally, the output $\boldsymbol{m}$ denotes the cumulative Qsoil lag offering the most explanatory power in terms of AIC values of PCA regression models and is thus a proxy for vegetation memory in terms of Qsoil with $\boldsymbol{n}$ identifying the same for air temperature.

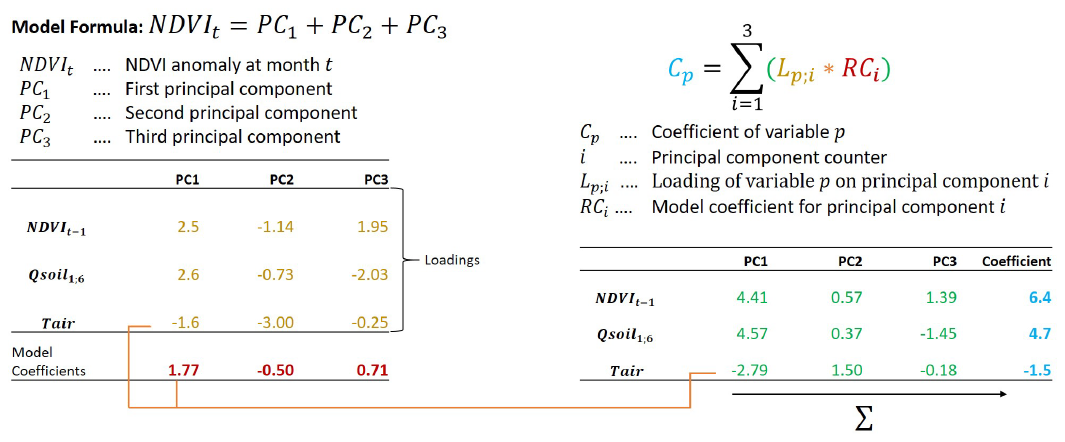

Figure 7 - PCA Regression Coefficients - Theoretical back-calculation of regression coefficients from PCA regression coefficients as lined out by Zuur et al., 2007. The data presented here (PCA loadings, PCA model coefficients, and final variable coefficients) represents a single pixel in the Iberian dryland region.

Interpretation

The duality of memory effects as intrinsic and extrinsic as proposed by Ogle et al., 2015 is embraced as follows:

1. **Intrinsic memory** is identified as coefficients of *NDVI[t−1]* (De Keersmaecker et al., 2015; Liu et al., 2018).
2. **Extrinsic memory** effects are implemented via ERA5 variables.
   a. *Tair* is implemented as cummulative lags ranging from 0 (instanteneous response) to 12 month-lags.
   b. *Qsoil1* - *Qsoil4* effects are implemented as cumulative lag effects ranging from instantaneous impacts to lags on annual time windows (Vicente-Serrano et al., 2013).

See Table 1 for an interpretation of the model coefficients within this context.

Table 1 - Interpretation of Memory Model Coefficients - Biological Interpretation of Vegetation Response Coefficients.

|  | **Magnitude** | **Sign** |
| --- | --- | --- |
| $\beta_{t-1}$ | Absolute values depict the speed at which systems return to equilibrium/pre disturbance state.  Large absolute values are assumed to represent low resilience (i.e. slow return) (De Keersmaecker et al., 2015). | *Positive* - NDVI anomalies resemble previous ones. NDVI anomalies gradually diminish.  *Negative* - NDVI anomalies resemble previous ones, but with the opposite sign. The return to pre-disturbance is characterised by oscillations. |
| $\beta_{Qsoil}$ | Absolute values depict the resistance to anomalies in Qsoil.  Large absolute values indicate low resistance (i.e. strong vegetation responses) to Qsoil anomalies. | *Positive -* Wetter conditions than average induce positive NDVI anomalies; drier soil conditions induce negative NDVI anomalies.  *Negative -* Drier conditions than average induce positive NDVI anomalies; wetter soil conditions induce negative NDVI anomalies. |
| $\beta_{Tair}$ | Absolute values depict the resistance to anomalies in air temperature.  Large absolute values indicate low resistance (i.e. strong vegetation responses) to air temperature anomalies. | *Positive -* Warmer temperatures than average induce positive NDVI anomalies; colder air temperature induces negative NDVI anomalies.  *Negative -* Colder temperatures than average induce positive NDVI anomalies; warmer air temperature induces negative NDVI anomalies. |

Variance Partitioning

Workflow

Variance partitioning for identification of vegetation memory component importance was carried out using the methodology presented by Real et al., 2003. First, we applied the full regression model $Z_{NDVI}=NDVI_{t-1}+Qsoil_{k;m}+ {Tair}_{n}$ and obtain $R^{2}$ (the coefficient of determination). This is $R_{Full}^{2}$ and equal to all explained variance. Unexplained variance can then be calculated as $1-R_{Full}^{2}$. Thereafter, we obtained $R^{2}$ of $Z_{NDVI}=NDVI_{t-1}$. This is $R_{NDVI_{t-1}}^{2}$ and equal to all variance explained by *NDVI[t−1]*. We then obtained $R^{2}$ of $Z_{NDVI}=Qsoil_{k;m}$. This is $R_{Qsoil_{k;m}}^{2}$ and equal to all variance explained by $Qsoil_{k;m}$ followed by $R_{Tair}^{2}$ as $Z_{NDVI}={Tair}_{n}$. Further shared variances are computed in the same manner and processed via the varPart function of the modEva package in R.

Results

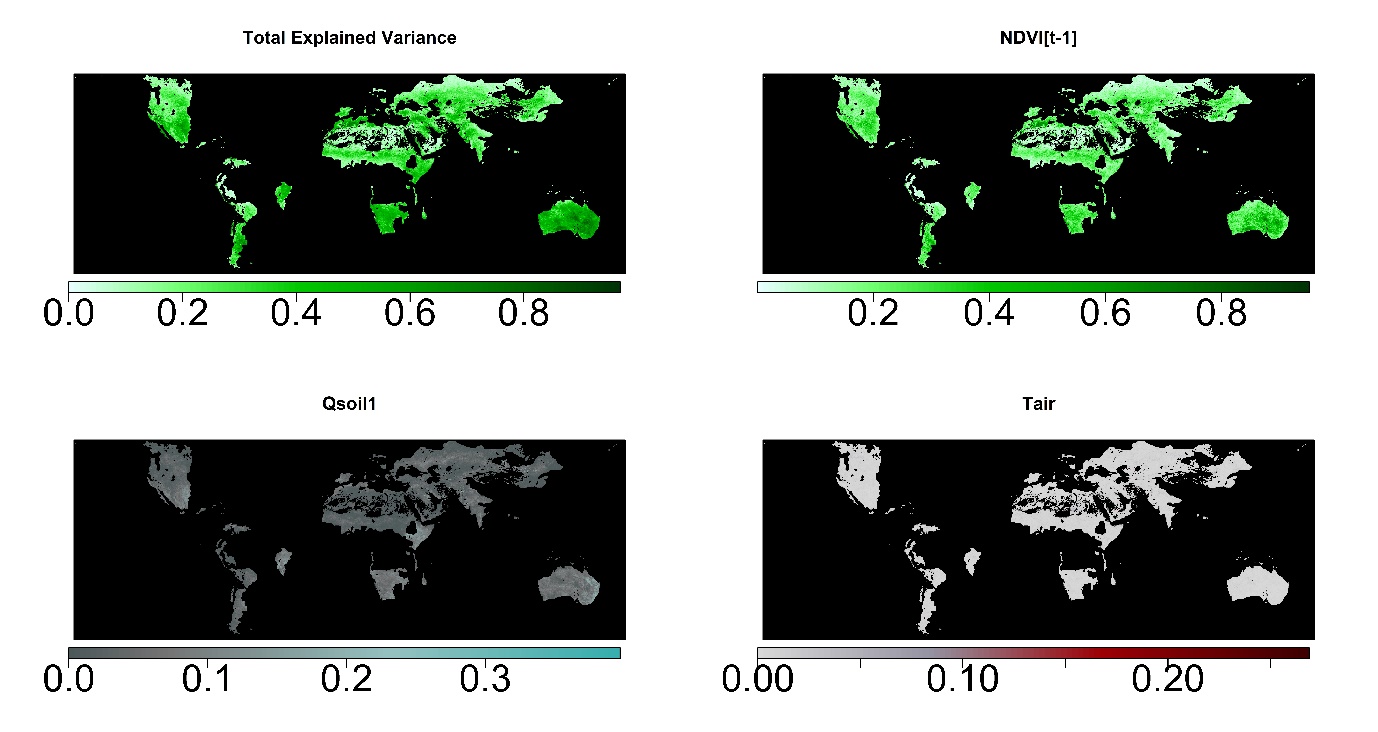

Figure 8 – Global Dryland Vegetation Memory Variance Partitioning 1 – Variance partitioning via partial regressions shows that our vegetation memory models perform well at explaining NDVI anomalies across global drylands with most variance explained by a single model component being that of intrinsic vegetation memory (*NDVI[t-1]*) followed by soil moisture (*Qsoil1*) vegetation memory. Variance explained by air temperature vegetation memory (Tair) is low and doesn’t show clear spatial patterns.

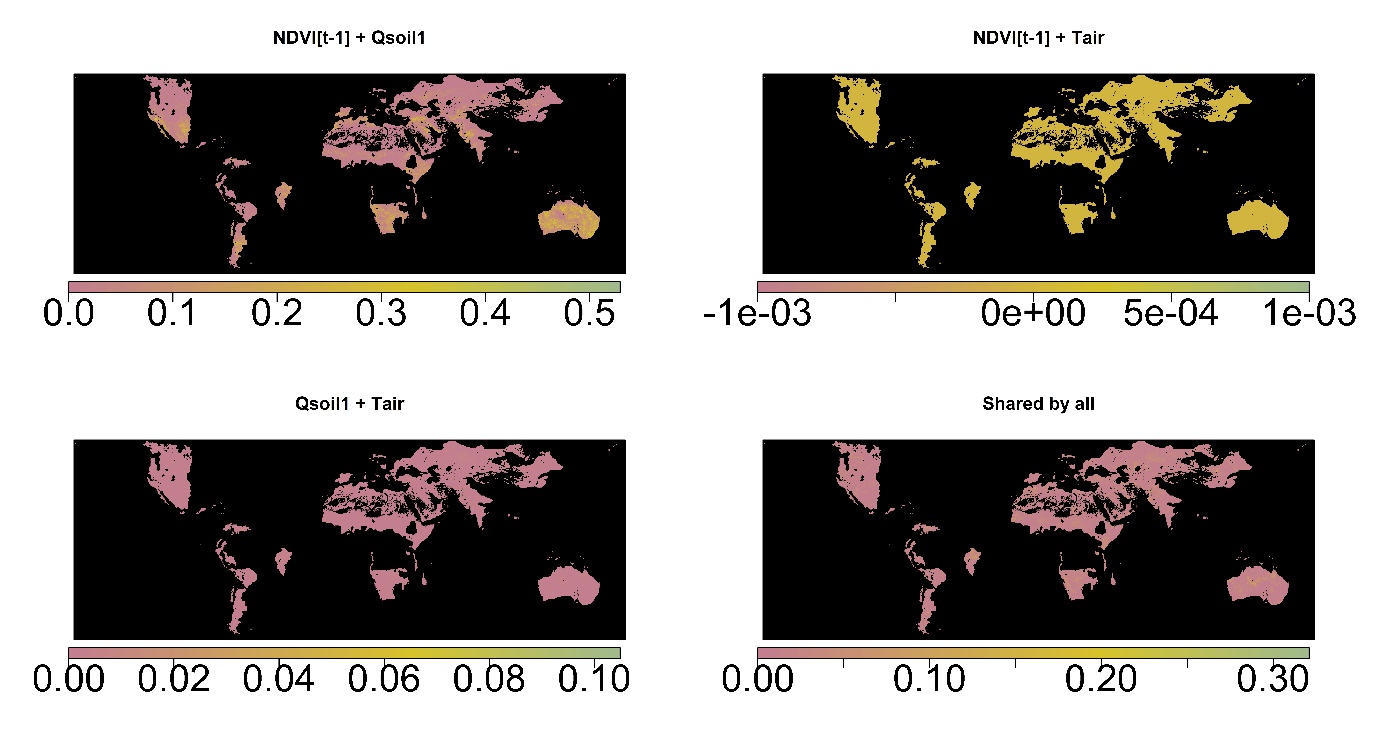

Figure 9 – Global Dryland Vegetation Memory Variance Partitioning 2 – Variance partitioning via partial regressions of shared variance in our vegetation memory models across global drylands shows that a lot of variance is shared by intrinsic memory and soil moisture memory (*NDVI[t-1] + Qsoil1*), while almost no variance in our models is shared between *Tair* and other variables.
